# The TrkA agonist gambogic amide augments skeletal adaptation to mechanical loading through actions on sensory nerves and osteoblasts

**DOI:** 10.1101/2020.08.28.272740

**Authors:** Gabriella Fioravanti, Phuong Q. Hua, Ryan E. Tomlinson

## Abstract

The periosteal and endosteal surfaces of mature bone are densely innervated by sensory nerves expressing TrkA, the high-affinity receptor for nerve growth factor (NGF). In previous work, we demonstrated that administration of exogenous NGF significantly increased load-induced bone formation through the activation of Wnt signaling. However, the translational potential of NGF is limited by the induction of substantial mechanical and thermal hyperalgesia in mice and humans. Here, we tested the effect of gambogic amide (GA), a recently identified robust small molecule agonist for TrkA, on hyperalgesia and load-induced bone formation. Behavioral analysis was used to assess pain up to one week after axial forelimb compression. Contrary to our expectations, GA treatment was not associated with diminished use of the loaded forelimb or sensitivity to thermal stimulus. Furthermore, dynamic histomorphometry revealed a significant increase in relative periosteal bone formation rate as compared to vehicle treatment. Additionally, we found that GA treatment was associated with an increase in the number of osteoblasts per bone surface in loaded limbs as well as a significant upregulation of *Wnt1, Wnt7b*, and *Ngf* in loaded bones. To determine if these effects were exclusively mediated by NGF-TrkA signaling in sensory nerves, we cultured MC3T3-E1 cells for 7 or 14 days in osteogenic differentiation media containing NGF (50 ng/mL), GA (5, 50, or 500 nM), or vehicle (DMSO). After 7 days of culture, we observed increases in osteoblastic differentiation markers *Runx2, Bglap2*, and *Sp7* in response to GA, whereas treatment with NGF was not different than vehicle. Only cells treated with the highest dose of GA (500 nM) had significantly impaired cell proliferation. In conclusion, our study indicates GA may be useful for augmenting skeletal adaptation to mechanical forces without inducing hyperalgesia through actions on both sensory nerves and osteoblasts.

## INTRODUCTION

The mammalian skeleton is highly responsive to mechanical stimuli^1^. Through a process known as mechanotransduction, bone cells sense and convert mechanical cues into biochemical signals, which subsequently direct and mediate both anabolic and catabolic processes. The signaling mechanisms that mediate load-induced bone formation have been studied extensively using a variety of experimental models^1,2^. Recent work from our lab and others has observed significant upregulation of nerve growth factor (NGF) in bone following both forelimb and tibial compression in mice^3–5^. Furthermore, we have shown that the inhibition of neurotrophic tyrosine kinase receptor 1 (TrkA), the high-affinity receptor for NGF expressed on the vast majority of sensory nerves in adult bone^6,7^, significantly diminished load-induced bone formation; on the other hand, administration of exogenous NGF significantly increased bone formation following loading^4^. In total, these experiments established the therapeutic potential of leveraging NGF-TrkA signaling to improve the anabolic response of the skeleton to mechanical load.

Unfortunately, administration of NGF is known to induce long-lasting mechanical and thermal hyperalgesia, as previously reported in both mice and humans^8–11^. Indeed, these painful side effects ultimately ended the promising clinical trials of recombinant human NGF to treat diabetes- and HIV-induced neuropathies^10^,^12–15^. In addition to causing unwanted side effects, NGF is an unlikely candidate to use as an anabolic bone agent due to the inherent drawbacks of using polypeptides as drugs, including poor stability and bioavailability^16^. As a result, leveraging NGF-TrkA signaling therapeutically to increase bone formation in response to load, and thereby decreasing the risk of fatigue injury, will require an alternative method for stimulating this signaling pathway in bone.

Recent work has endeavored to characterize small and stable molecules that selectively bind to TrkA^17–20^. Of particular note is gambogic amide (GA), a small molecule (627.8 Da) uncovered in a cell-based chemical genetic screen designed to identify TrkA agonists^18^. Similar to NGF, GA was found to significantly inhibit glutamate-induced neuronal cell death and induce robust neurite outgrowth in PC12 cells^18^. However, rather than inducing the dimerization of TrkA by binding to the extracellular ligand-binding region, GA appears to bind to the intracellular juxtamembrane domain of TrkA and facilitates NGF activity through allosteric activation of TrkA^21^. As a result, GA induces lower magnitude but longer lasting TrkA phosphorylation^20^. Importantly, GA is inexpensive, well-tolerated in vivo, and readily available in large quantities.

In this study, we investigated the effect of GA on mice subjected to axial forelimb compression as well as MC3T3-E1 cells in culture. Our overall hypothesis was that administration of GA would increase NGF-TrkA signaling in bone following mechanical loading, leading to increases in load-induced bone formation and anabolic signaling, without the induction of marked thermal or mechanical hyperalgesia. Furthermore, we hypothesized that GA would not affect osteoblastic cells directly as they generally do not express TrkA, the high affinity receptor for NGF and target of GA. The results from our study reveal novel actions of GA on both sensory nerves and osteoblasts.

## METHODS

### Mice

All procedures were approved by the Institutional Animal Care and Use Committee of Thomas Jefferson University (#02204). Adult C57BL/6J mice (Jackson Laboratory #000664) were used for all studies.

### Mechanical Loading

Gambogic amide (0.4 mg/kg) or vehicle (DMSO) was administered one hour prior to loading. In experiments with multiple days of loading, GA or vehicle was only administered on day 0. Immediately before loading, mice were then anesthetized using isoflurane gas (2-3%) and received buprenorphine (0.12 mg/kg, IP). Next, the right forelimb was axially compressed in specially designed fixtures using a material testing system (TA Systems Electroforce 3200). A 0.3 N preload was applied, followed by a cyclic rest-inserted trapezoidal waveform with a peak force of 3 N at 2 Hz for 100 cycles. The left forelimb was not loaded and served as a contralateral control. Mice were allowed unrestricted cage activity after loading.

### Histomorphometry

Bone formation rates were quantified by dynamic histomorphometry using undecalcified sections from the mid-diaphysis of loaded and non-loaded forelimbs. Mice were given intraperitoneal injections of calcein (10 mg/kg; Sigma C0875) and alizarin red (30 mg/kg; Sigma A3882) at day 3 and 8, respectively. Forelimbs were harvested at day 10, fixed in 10% neutral buffered formalin for 16-24 hours, and embedded in polymethylmethacrylate. Samples were sectioned at 100 μm using a low-speed saw (Isomet 1000) and mounted on glass slides with Eukitt mounting medium (Sigma 03989). After drying, sections were then polished to 50 μm and imagined using fluorescence microscopy (Nikon Eclipse E800). Images were analyzed for endosteal (Es) and periosteal (Ps) mineralizing surface (MS/BS), mineral apposition rate (MAR), and bone formation rate (BFR/BS). For analysis by static histomorphometry, sections were further polished, then stained with 50°C preheated Sanderson’s Rapid Bone Stain (Dorn & Hart Microedge S-SRBS1) for 30 seconds. Next, sections were then counterstained with room temperature Acid Fuchsin for 10 seconds and quickly dehydrated in 100% ethanol. Sections were imaged using bright-field microscopy and analyzed for the number of osteoblasts per bone surface and osteocytes per bone area.

### Mechanical and Thermal Sensitivity

Analyses were performed one day before the first bout of loading and then 1, 4, and 7 days following the final bout of loading. First, forelimb asymmetry testing was used to assess overall mechanical sensitivity of the loaded limb relative to the non-loaded limb. As in previous studies^22^, mice recorded for 5 minutes after being placed inside a clear cylindrical tube. Mirrors were positioned to allow visual inspection of the entire tube at once. Each incidence of vertical exploration was scored, with a score of 1 given for the right (loaded) forepaw, 0.5 for both forepaws, and 0 for only the left (non-loaded) forepaw. Following this, thermal sensitivity was assessed by measuring the response time of each mouse to a hotplate maintained at 55°C, as in previous work^4^. Mice were immediately removed from the hot plate following a paw lick, paw flick, jump, or after 30 seconds has elapsed without a response. Quantification was performed after the test using a video recording.

### Osteoblast Culture

MC3T3-E1 Subclone 4 (ATCC CRL-2593) cells were recovered from liquid nitrogen and cultured to confluency in α-MEM (Corning, Mediatech, Inc) supplemented with 10% fetal bovine serum and 1% penicillin/streptomycin in a 37 °C humidified incubator at 5% CO2. Differentiation was induced by addition of 10mM β-glycerol phosphate and 50 μg/ml ascorbic acid to the media after plating. For alizarin red staining and mRNA collection, cells were placed into six-well plates at a density of 50,000 cells/well. For apoptosis and proliferation assays, cells were seeded in 96-well plates at a density of 20,000 cells/well or 5,000 cells/well respectively. Cells were treated with either vehicle (DMSO), 50 ng/mL nerve growth factor (Envigo NGF 2.5S), or 5nM, 50 nM, or 500nM GA (Enzo Life Sciences BML-N159-0001) continuously over the course of each experiment.

### In vitro assays

Mineralization was quantified following incubation for 3, 7, 14, or 21 days in osteogenic media. Alizarin red staining was performed using prepared reagents, according to the manufacturer’s instructions (ScienCell, ARed-Q). Stain absorbed by cells was quantified by reading the absorbance of the cell using a plate reader (Tecan M1000) at 405 nm. Here, absorbance directly correlates to the total alizarin red staining in each well. Cell proliferation was determined using CellTiter 96 AQueous One Solution Cell Proliferation Assay kit (Promega Corporation) as per manufacturer’s instructions and quantified by absorbance at 490nm. Cell apoptosis was determined using HT TiterTACS Assay Kit (Trevigen) as per manufacturer’s instructions and quantified by absorbance at 450nm.

### qRT-PCR

Expression of osteoblastic gene markers in MC3T3 cells was quantified by qRT-PCR after 3, 7, 14, and 21 days of osteogenic differentiation. RNA (0.5 ug) isolated using TRIzol (Life Technologies) was reverse transcribed using iScript cDNA Synthesis Kit (Bio-Rad), then cDNA (1.6 uL) was amplified under standard PCR conditions using PowerUp SYBR (Bio-Rad). Similarly, gene expression in forelimbs either 3 or 24 hours after a single bout of mechanical loading was quantified by qRT-PCR. After harvesting forelimbs, the proximal and distal ends of the bone were cut off and the marrow was removed by brief centrifugation at 13000g before placing into TRIzol (Ambion). After pulverization in liquid nitrogen (SpexMill 6750), total RNA was extracted from both forelimbs using TRIzol. Similar to the above, RNA (0.5μg) was reversely transcribed using iScript Reverse Transcription Supermix (Bio-Rad), and cDNA (1.6 μL) was amplified under standard PCR conditions using PowerUp SYBR (Bio-Rad). For all samples, cDNA was amplified in triplicate and normalized to GAPDH expression. Fold changes of loaded limbs vs. non-loaded limbs were computed using the ddCT. Primer sequences were designed using Primer-BLAST (NCBI) and are available in Table 1.

**Table 1.**
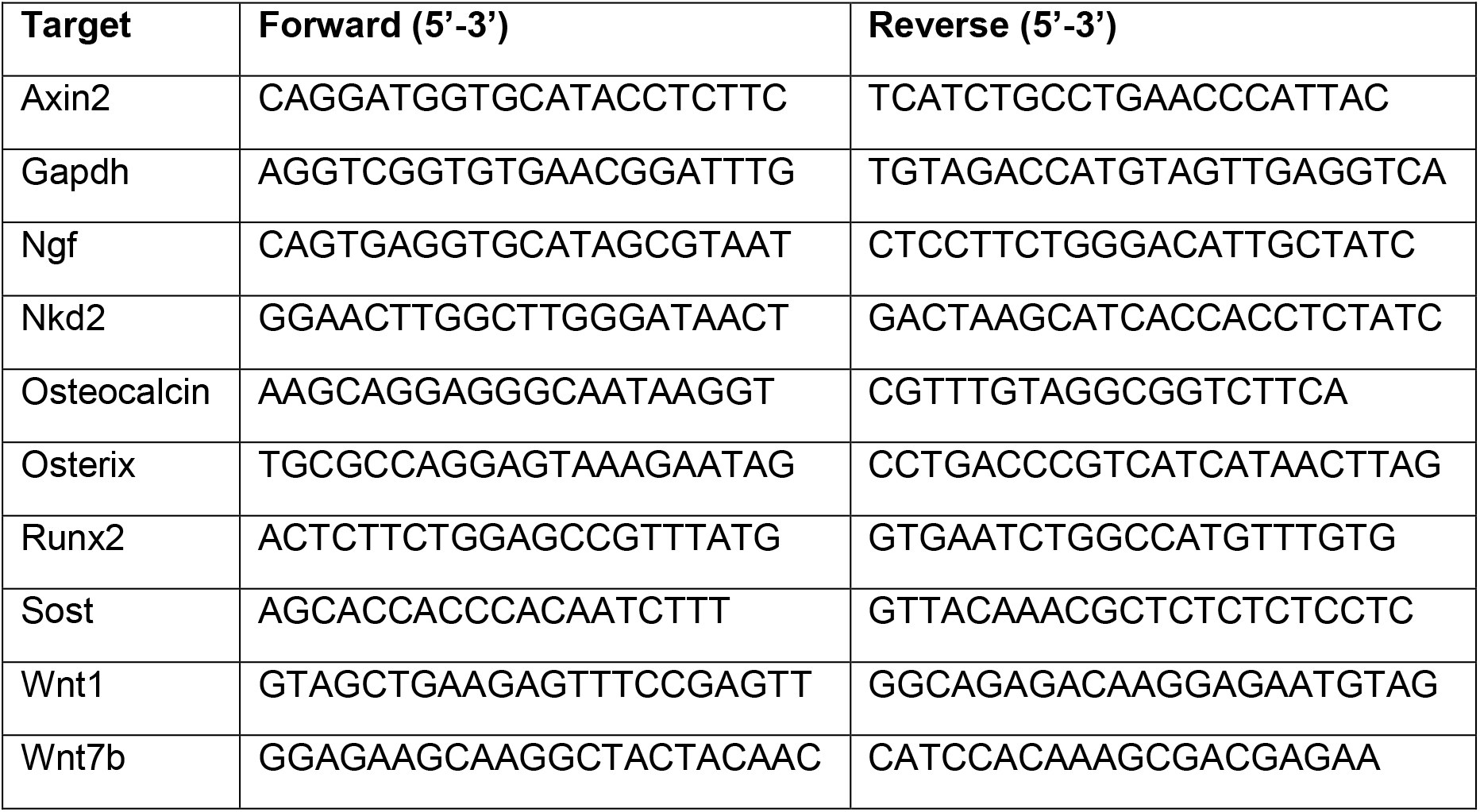
Oligonucleotide primers used for qRT-PCR.

### Statistics

All results are presented as mean ± SE. Statistical analysis were performed on Prism 8 (GraphPad) using unpaired, two-tailed Student’s t-test when comparing two groups or a two-way ANOVA test when comparing more than two groups. Statistical significance level was determined when p < 0.05, denoted with * in graphical figures.

## RESULTS

### Gambogic amide increases load-induced bone formation

To determine if administration of gambogic amide (GA) increases load-induced bone formation, adult C57BL6/J mice were subjected to three consecutive bouts of axial forelimb compression designed to produce lamellar bone formation. Either GA (0.4 mg/kg) or vehicle (DMSO) was injected 1 hour before the first bout of loading. Calcein and alizarin red bone formation labels administered 3 and 8 days after the first bout of loading were visualized in PMMA-embedded sections (Fig. 1A,B) and quantified using dynamic histomorphometry (Table 2). As expected, forelimb loading induced a robust periosteal bone formation response in both GA and vehicle treated mice. Whereas GA did not significantly increase relative (loaded – non-loaded) periosteal mineralizing surface (Fig. 1C), administration of GA was associated with a significant increase (+63%) in relative periosteal mineral apposition rate in response to loading (Fig. 1D). As a result, treatment with GA was associated with a significant increase (+63%) in relative periosteal bone formation rate (Fig. 1E). Sections stained with Sanderson’s Rapid Bone Stain (Fig. 2A) revealed no effect of GA on osteocyte number in either loaded or non-loaded limbs (Fig. 2B), but a significant increase in the number of osteoblasts per millimeter of bone surface in loaded limbs (Fig. 2C).

**Table 2.** Gambogic amide increases load-induced bone formation. Values are presented as mean ± standard deviation. * p < 0.05 vs. Non-Loaded. n=7 per group.

| GA |  |  |  |  |  |  |
| --- | --- | --- | --- | --- | --- | --- |
| | PS.MS/BS<br>($\mu\text{m}/\mu\text{m}$) | Es.MS/BS<br>($\mu\text{m}/\mu\text{m}$) | Ps.MAR<br>($\mu\text{m}/\text{day}$) | Es.MAR<br>($\mu\text{m}/\text{day}$) | Ps.BFR/BS<br>($\mu\text{m}^3/\mu\text{m}^2/\text{day}$) | Es.BFR/BS<br>($\mu\text{m}^3/\mu\text{m}^2/\text{day}$) |
| **Loaded** | 0.41 ±<br>0.12 * | 0.80 ±<br>0.09 * | 1.16 ±<br>0.66 * | 1.35 ±<br>0.51 * | 0.43 ± 0.16 * | 1.12 ± 0.47 * |
| **Non-Loaded** | 0.22 ±<br>0.15 | 0.64 ±<br>0.12 | 0.29 ±<br>0.13 | 0.67 ±<br>0.26 | 0.08 ± 0.07 | 0.41 ± 0.16 |
| Vehicle |  |  |  |  |  |  |
| | PS.MS/BS<br>($\mu\text{m}/\mu\text{m}$) | Es.MS/BS<br>($\mu\text{m}/\mu\text{m}$) | Ps.MAR<br>($\mu\text{m}/\text{day}$) | Es.MAR<br>($\mu\text{m}/\text{day}$) | Ps.BFR/BS<br>($\mu\text{m}^3/\mu\text{m}^2/\text{day}$) | Es.BFR/BS<br>($\mu\text{m}^3/\mu\text{m}^2/\text{day}$) |
| **Loaded** | 0.35 ±<br>0.12 * | 0.72 ±<br>0.22 | 0.68 ±<br>0.18 * | 1.06 ±<br>0.38 * | 0.25 ± 0.12 * | 0.76 ± 0.38 * |
| **Non-Loaded** | 0.19 ±<br>0.12 | 0.52 ±<br>0.23 | 0.15 ±<br>0.08 | 0.46 ±<br>0.17 | 0.03 ± 0.03 | 0.25 ± 0.12 |

**Figure 1.**
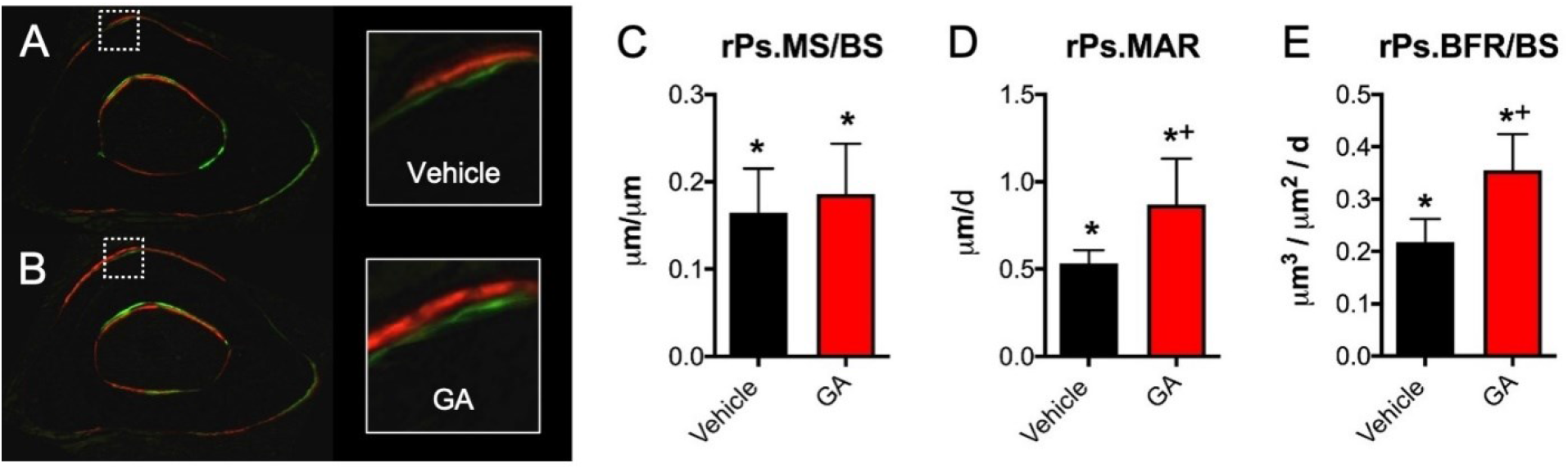
GA increases periosteal bone formation following axial forelimb compression. A,B) Calcein (green) and alizarin red (red) fluorescent bone formation markers were injected following 3 days of axial forelimb compression. Relative (loaded – non-loaded) periosteal bone formation parameters C) rPs.MS/BS, D) rPs.MAR, and E) rPS.BFS/BS were quantified. *p < 0.05 vs. non-loaded, +p < 0.05 vs. vehicle.

**Figure 2.**
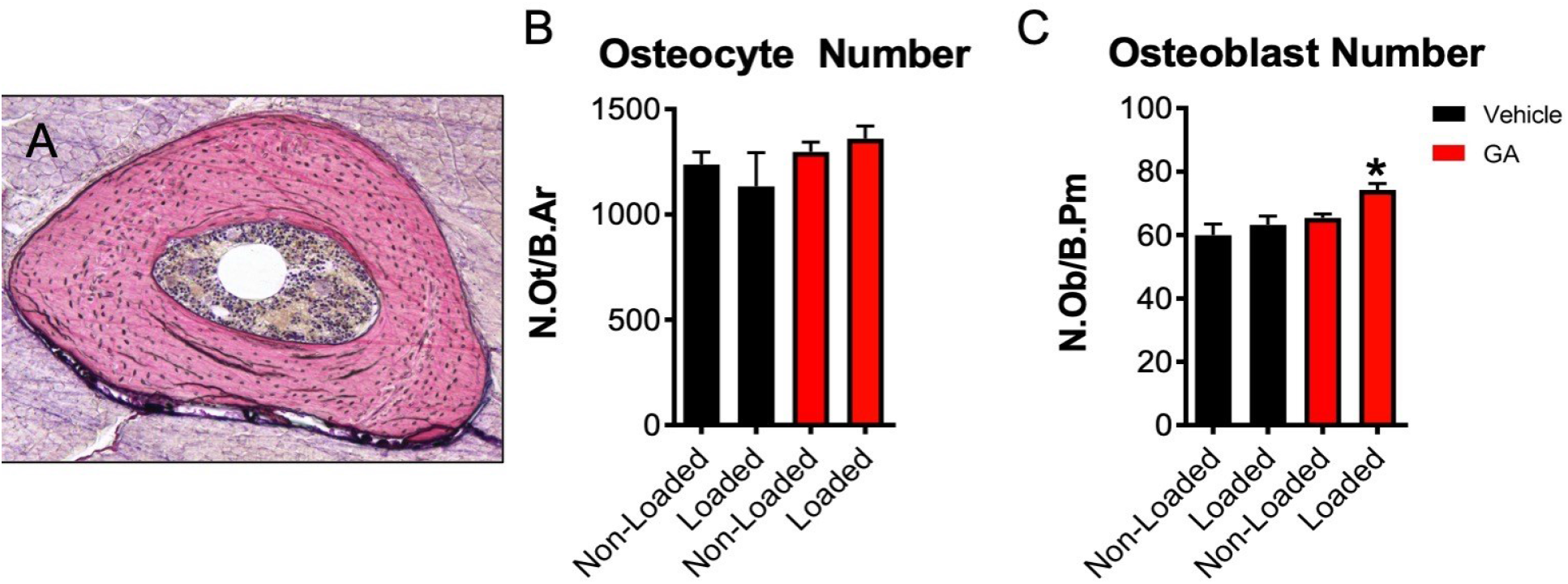
GA increased osteoblast number in loaded limbs. A) Sections were stained with SRBS and Acid Fuchsin for quantification of B) osteocytes/bone area (N.Ot/B.Ar) and C) osteoblasts/bone surface (N.Ob/B.Pm). * p < 0.05 vs. non-loaded.

### Gambogic amide does not induce mechanical or thermal hyperalgesia

To determine if GA induced the painful side effects reported following administration of NGF, we observed forelimb asymmetry and hotplate response one day before the first bout of loading (baseline) as well as 1, 4, and 7 days following the final bout of loading. On the 1^st^ day following axial forelimb compression, we observed significantly less usage of the loaded limbs of vehicle treated mice (−9% vs. baseline), whereas GA treated mice displayed no significant differences in limb preference (Fig. 3A). At the 4^th^ day and 7^th^ day after loading, there were no significant differences between treatment groups or loading conditions. At these same time points, we quantified thermal sensitivity using standard hotplate analysis (Fig. 3B). Similar to forelimb asymmetry testing, we observed that GA treated mice, but not vehicle treated mice, took significantly longer to respond to the hot plate as compared to baseline 1 day after loading. However, there were no significant differences between treatment groups or loading conditions at day 4 or 7. In total, these data indicate that GA does not induce hyperalgesia in mice, particularly following osteogenic mechanical loading.

**Figure 3.**
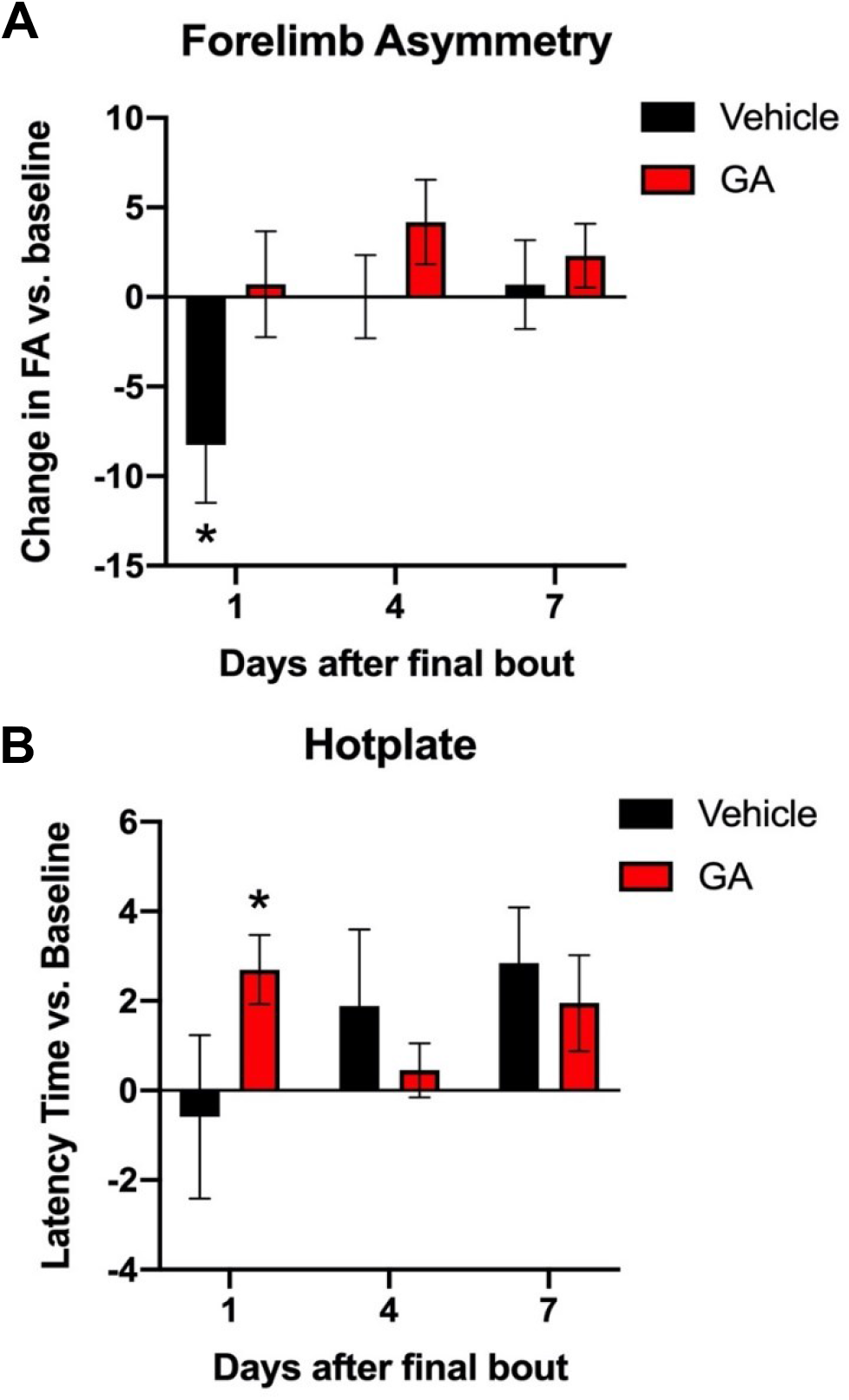
GA decreased mechanical and thermal sensitivity. A) Forelimb usage after loading was assessed by quantitative of forelimb asymmetry, where a decreased score indicates less usage of the loaded limb. B) Thermal sensitivity was assessed by latency time after hotplate challenge, where an increased score indicates less sensitivity to thermal stimulus. * p < 0.05 vs. baseline.

**Figure 4.**
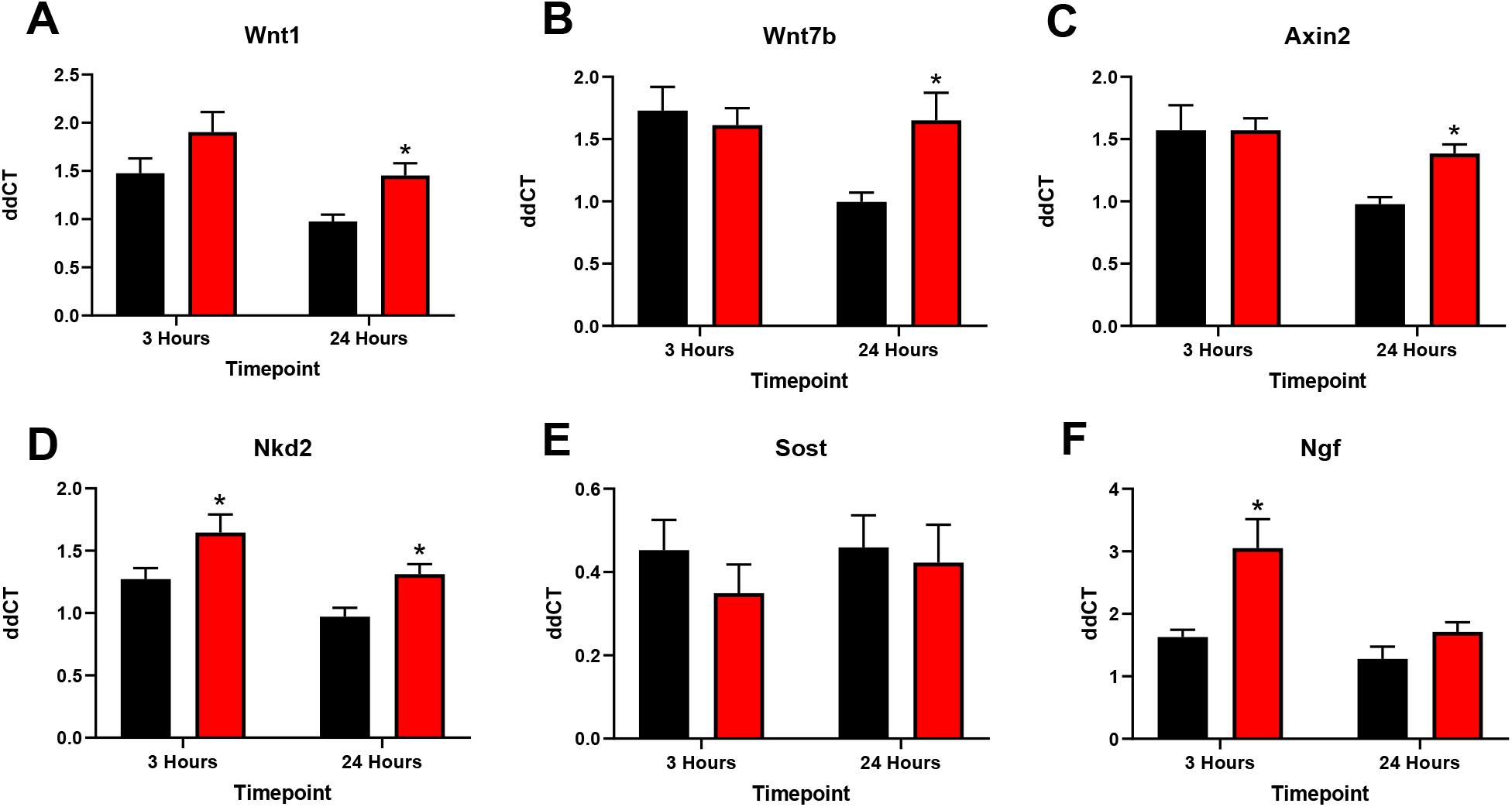
GA upregulates Wnt signaling in loaded bone. Relative expression levels of mRNA in the forelimb 3 and 24 hours after one bout of loading (100 cycles of 3N sinusoidal waveform at 2 Hz rest inserted) for A) Wnt1 B) Wnt7b C) Axin2 D) Nkd2 E) Sost and F) NGF * p < 0.05 vs. vehicle; n=7-10 per group

### Gambogic amide increases osteogenic gene transcription following loading

To determine the specific effects of GA on osteogenic gene expression, we harvested mRNA from the central third of loaded and non-loaded ulna after either 3 or 24 hours following a single bout of loading for analysis by qRT-PCR. Here, mice were injected with either GA (0.4 mg/kg) or vehicle (DMSO) 1 hour prior to loading. Similar to the effects of exogenous NGF^4^, GA significantly increased the expression of Wnt ligands (*Wnt1, Wnt7b*) and Wnt target genes (*Axin2, Nkd2*) in bone 24 hours after loading; only *Nkd2* was significantly increased after 3 hours. Consistent with previous studies, *Ngf* was significantly upregulated by loading alone after 3 hours, and GA treatment significantly increased the expression of *Ngf* at 3 hours as compared to vehicle. Finally, *Sost* was significantly downregulated by loading, but there was no additional downregulation of *Sost* associated with GA at either 3 or 24 hours. In total, these results indicate that GA increases gene transcription associated with load-induced bone formation, potentially by amplifying the expression of *Ngf* in mature osteoblasts.

### Gambogic amide increased osteoblastic differentiation markers *in vitro*

Since our *in vivo* data suggested that GA may directly affect osteoblasts, we performed additional *in vitro* qRT-PCR using mRNA harvested from MC3T3-E1 cells that were incubated in osteoblastic differentiation media containing GA (5-500 nM), NGF (50 ng/mL), or vehicle (DMSO) control. Consistent with previous studies, we were unable to detect TrkA expression at any point during 21 days of differentiation (S. Fig 1), and there were no significant effects of NGF. In contrast, media containing GA induced the significant upregulation of the osteoblastic differentiation markers *Runx2, Bglap2*, and *Sp7* in a dose-dependent manner (Fig. 5). Consistent with our *in vivo* findings, we observed that administration of 50 nM of GA upregulated expression of *Ngf* at both Day 3 and Day 7. In total, these results indicate that GA acts directly on osteoblasts to increase osteoblastic differentiation markers and upregulate *Ngf* expression in non-loaded conditions.

**Figure 5.**
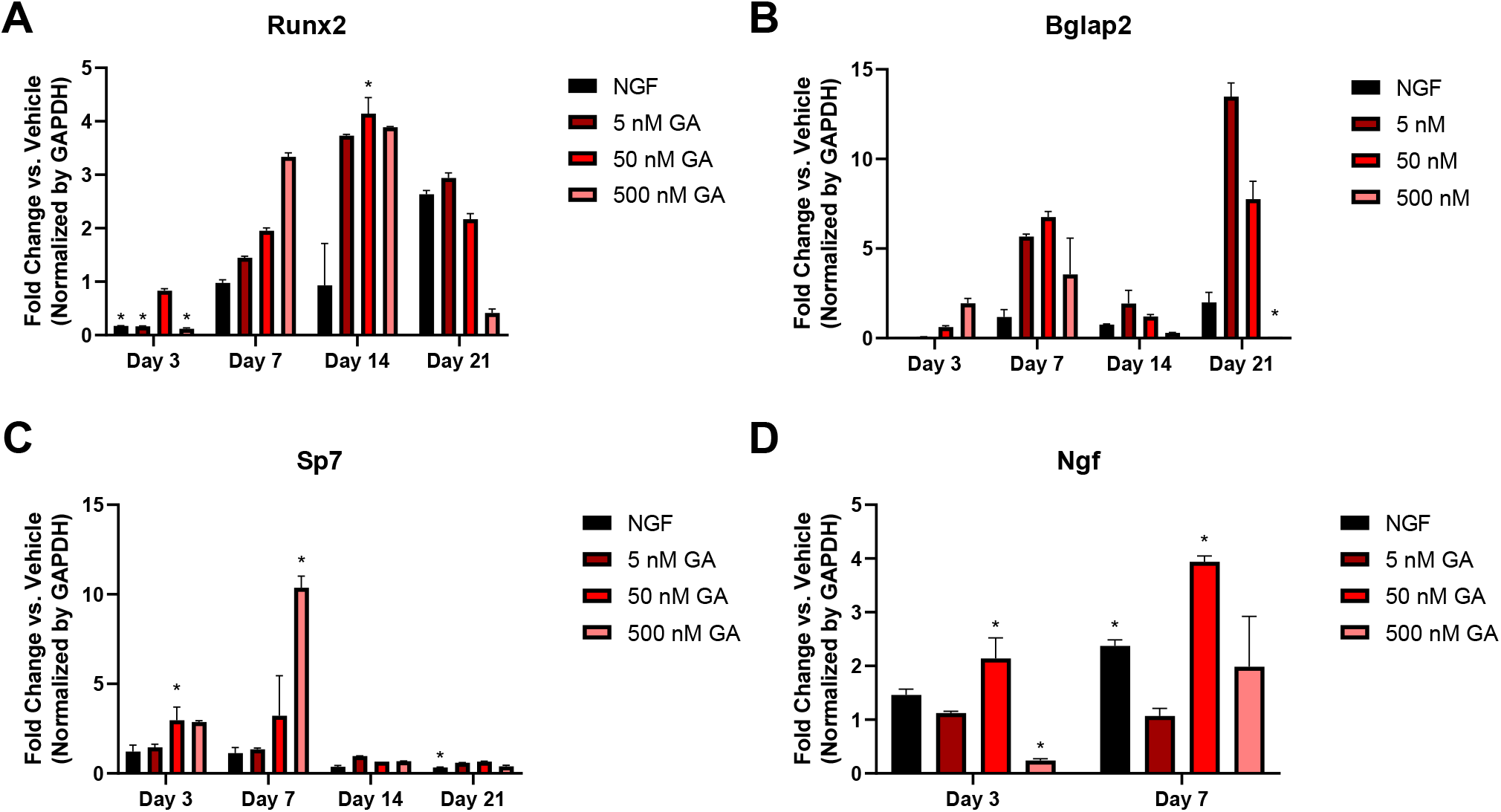
GA increases osteoblastic differentiation markers in MC3T3 cells. Relative expression levels of mRNA after treatment of Vehicle (DMSO), NGF, or 5, 50, 500 nM GA. Fold change versus vehicle shown for A) Runx2, B) Bglap2, C) Sp7, and D) NGF spanning up to 3 weeks. * p < 0.05 vs. vehicle

### Gambogic amide effects on osteoblast proliferation, apoptosis, and mineralization *in vitro*

To further explore the effects on GA on osteoblasts, we performed a proliferation assay on MC3T3-E1 cells cultured in complete media containing GA (5-500 nM), NGF (50 ng/mL), or vehicle (DMSO) control for 72 hours. Although neither NGF nor the 5 or 50 nM concentration of GA affected the proliferation rate, cells treated with 500 nM of GA had significantly impaired cell proliferation (Fig. 6A). Next, we performed an assay to determine if GA induced apoptosis in osteoblasts. Here, MC3T3-E1 cells were cultured in complete media containing GA (5-500 nM), NGF (50 ng/mL), or vehicle (DMSO) control for 96 hours (Fig. 6B). Only the group treated with 50 nM of GA had significantly greater apoptosis than vehicle (+245%), and the group treated with 500 nM trended toward significance (+170%, P = 0.2957). Finally, we determined the effect of GA on mineralization by alizarin red staining. Here, MC3T3-E1 cells were cultured in osteogenic media containing GA (5-500 nM), NGF (50 ng/mL), or vehicle (DMSO) control for 21 hours. There were no significant differences in any treatment group after 3 or 7 days of culture (Fig. 6C,D). After 14 days, we observed a significant increase in alizarin red staining in the group treated with 5 nM GA as compared to vehicle (Fig. 6E), but there were no significant differences at Day 21 (Fig. 6F). In total, these results suggest that GA primarily acts through its activation of non-osteoblastic TrkA to affect bone formation, with moderate doses having limited effects on proliferation, apoptosis, and mineralization *in vitro*. Furthermore, these results confirm that NGF has no direct effects on osteoblasts, consistent with their lack of expression of TrkA.

**Figure 6.**
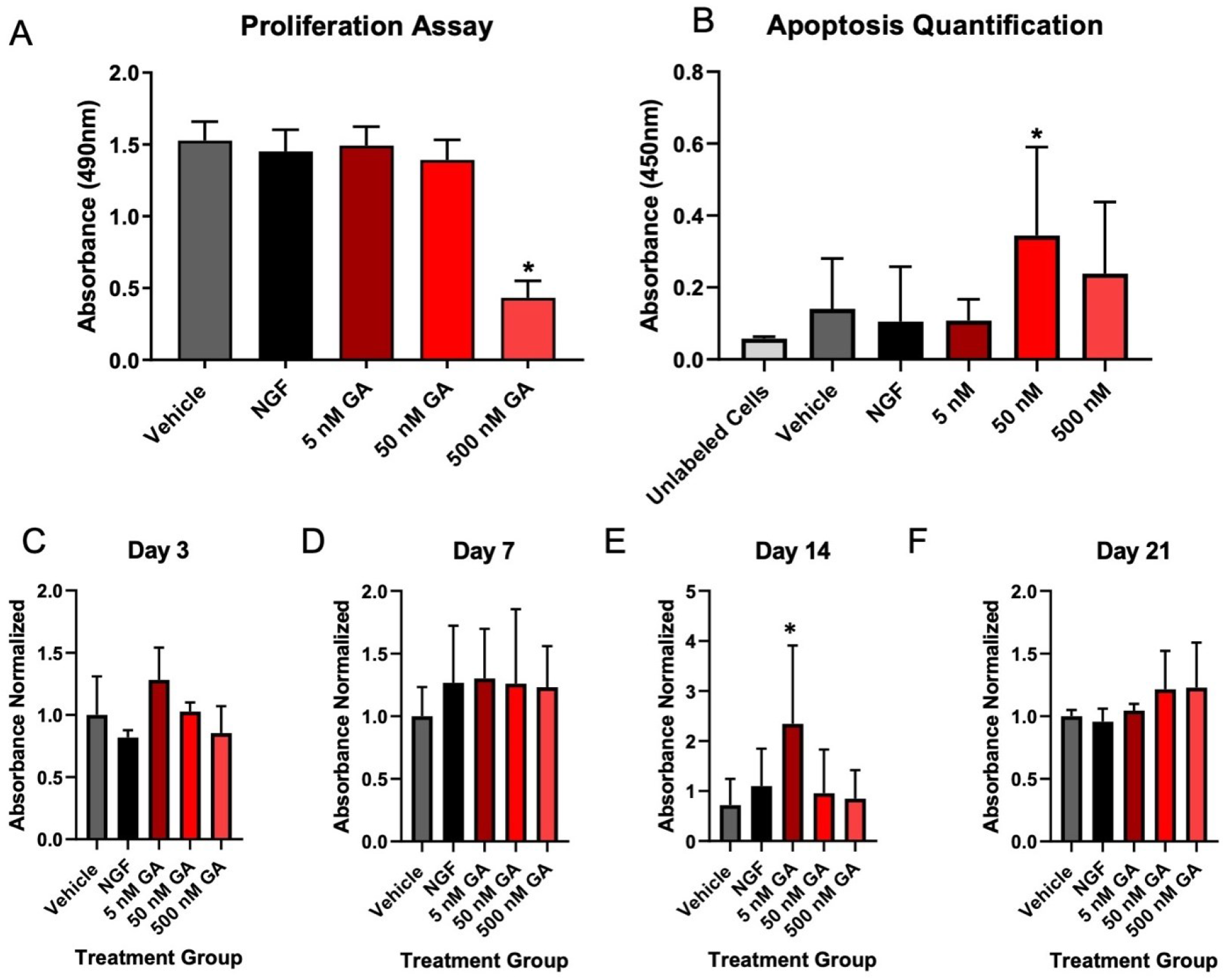
Gambogic amide mildly affects proliferation, apoptosis, and mineralization in MC3T3 cells at higher concentrations. A) Proliferation of cells treated with Vehicle (DMSO), NGF, or a low, moderate, or high dose of GA as quantified by 72-hour colorimetric MTS assay. B) Quantification of apoptosis of MC3T3 cells for the same 5 groups. C-F) Alizarin red staining was quantified in MC3T3 cells cultured for 3-21 days in osteogenic media. * p < 0.05 vs. vehicle

## DISCUSSION

Our main objective in this study was to examine the action of the TrkA agonist gambogic amide (GA) on load-induced bone formation and hyperalgesia in mice, with the long-term goal of utilizing this small molecule to increase bone mass in patients at risk for stress fracture without the negative side effects of NGF. The results from our study indicate that GA may be a potential novel therapeutic for increasing bone formation rate following loading. Importantly, we observed a significant increase in relative periosteal bone formation rate following axial forelimb compression that was not associated with increased thermal or mechanical sensitivity. Furthermore, we observed that GA appears to act on both TrkA expressing sensory nerves and osteoblasts, in contrast to the action of NGF which is confined only to T rkA expressing nerve axons in bone (Fig. 7). In summary, GA may be useful for leveraging NGF-TrkA signaling to increase skeletal adaptation to mechanical forces without inducing hyperalgesia.

**Figure 7.**
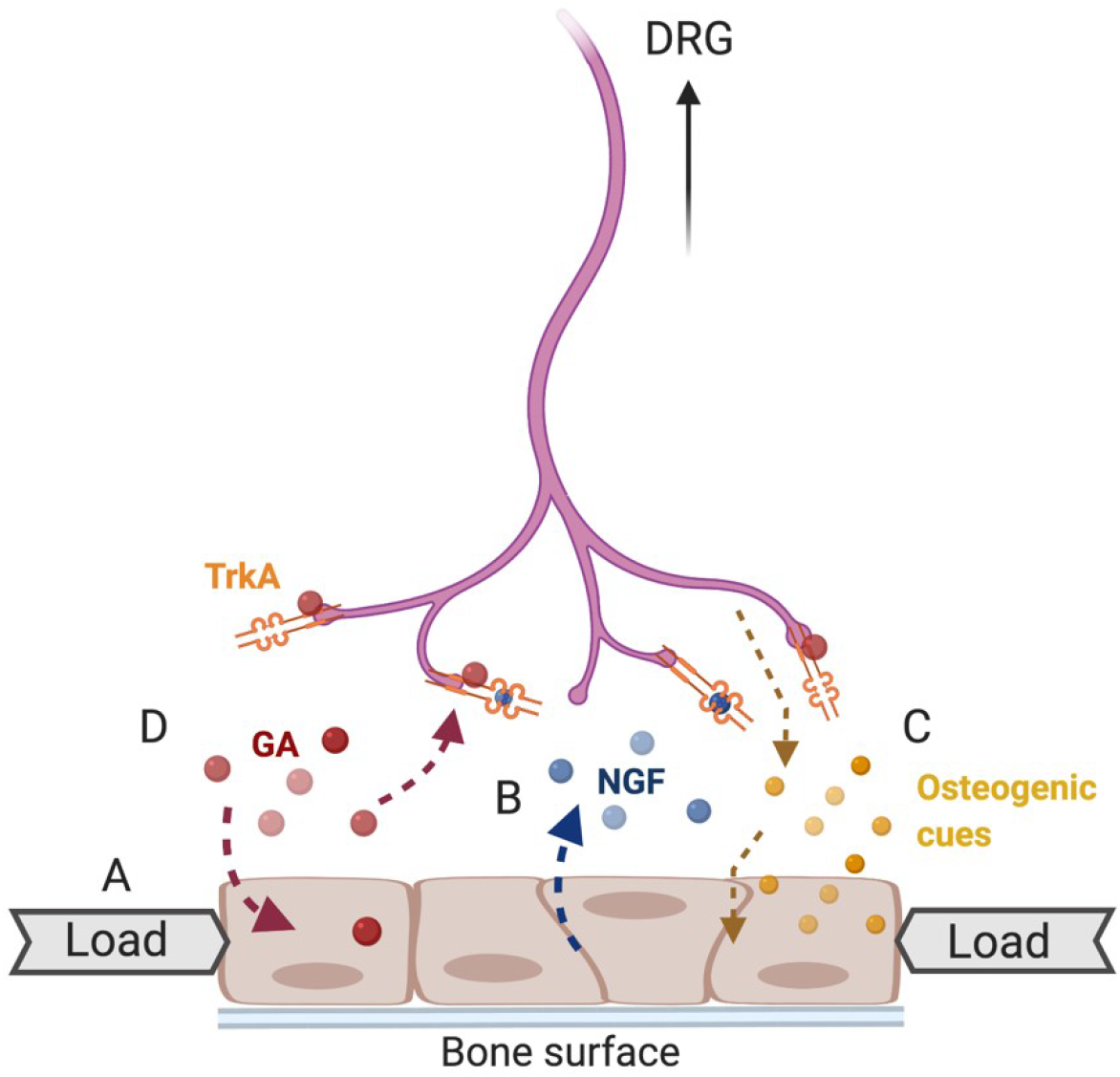
Schematic of GA action on load-induced bone formation. A) In response to mechanical loading, mature osteoblasts on the bone surface express B) NGF that initiates signaling in TrkA-expressing skeletal sensory nerves extending from the DRG to release C) osteogenic cues. D) GA can bind allosterically to TrkA receptors to facilitate NGF-TrkA signaling as well as act directly on osteoblasts to increase osteoblast differentiation and stimulate NGF expression. Created with BioRender.com.

We have previously shown that activation of NGF-TrkA signaling by a single administration of exogenous NGF (5 mg/kg BW) was sufficient to increase load-induced bone formation, but resulted in significant thermal hyperalgesia for at least 72 hours^4^. Here, we used forelimb asymmetry and hotplate testing to assay the response to GA following loading. Contrary to our expectation, GA treatment was not associated with diminished use of the loaded forelimb by forelimb asymmetry testing at any time point, whereas vehicle treated mice used their loaded limb less 1 day after the final bout of loading. Similarly, we observed less sensitivity to the hot plate in GA treated mice 1 day after the final bout of loading, but no significant differences in vehicle treated mice at any time point. Although unexpected, these results indicate that an effective dose of GA does not result in the potent hyperalgesia observed after administration of NGF. Moreover, our results are broadly consistent with previous work that found that a lower dose of GA (5 mM) directly injected to L4/L5 DRG by lumbar puncture induced only mild thermal hyperalgesia for 2 days and had no significant effect on mechanical hyperalgesia for at least 14 days^23^.

To determine the mechanism by which GA increased load-induced bone formation following axial forelimb compression, we assayed the Wnt/β-catenin signaling pathway, which must be activated for a normal anabolic response to mechanical loading in bone^24,25^. Our results indicate that the increased NGF-TrkA signaling induced by GA led to significant increases in Wnt ligands and target genes in bone in the first 24 hours after loading, consistent with previous studies examining the role of NGF following mechanical load^4^. Surprisingly, we also observed that expression of NGF itself was significantly upregulated by administration of GA in mice as well as in MC3T3-E1 cells cultured in GA-containing media. To our knowledge, this effect has not yet been reported. However, a previous group demonstrated that administration of GA upregulated TrkA protein and mRNA *in vitro* and *in vivo^20^*, consistent with previous studies showing that TrkA is a transcriptional target of NGF^26^. As a result, we acknowledge the possibility that part or all the effect of GA on bone is mediated by increased NGF expression from osteoblasts and/or osteocytes, rather than the action of GA itself on bone cells and sensory nerves. However, more study using mice deficient in NGF would be required to determine the extent to which the action of GA is dependent on NGF.

Nonetheless, to determine the direct effects of GA on osteoblasts, we performed a series of *in vitro* experiments using MC3T3-E1 osteoblasts. In previous work, we observed that neither mouse MSCs nor mouse calvarial osteoblasts expressed TrkA or responded to TrkA inhibition^27^. As a result, we hypothesized that NGF and GA would act primarily on sensory nerves in bone, rather than osteoblast-lineage cells. Consistent with this hypothesis and the lack of TrkA expression in these cells (S. Fig. 1), we observed no significant increases in osteoblast differentiation markers, proliferation, apoptosis, or mineralization in MC3T3-E1 cells cultured in osteogenic media containing NGF (50 ng/mL). In contrast, GA significantly increased osteoblast differentiation markers in a dose dependent manner, the highest concentrations of GA decreased proliferation and increased apoptosis, and mineralization was modestly increased. These results suggest that GA, but not NGF, can directly affect proliferating and mature osteoblasts in bone. Similar to our study, researchers found that treatment of MC3T3-E1 cells with NGF (10 to 400 pg/mL) did not affect proliferation^28^. However, in contrast to our assertion that NGF does not affect osteoblasts, they reported that NGF increased alkaline phosphatase activity and type 1 collagen production, and these effects could be blocked by an anti-NGF antibody. Similarly, another group reported that GA treatment increased expression of alkaline phosphatase, osteocalcin, and DMP-1 in Kusa O cells after 14 days of culture in osteogenic media, which is consistent with our results^29^. However, they attributed these effects to the induction of TrkA expression after differentiation of Kusa O cells in osteogenic media for 14 days^29^, a phenomenon that we did not observe in MC3T3-E1 cells. As a result, more study is necessary to clarify the effects of both GA and NGF on human osteoblasts before development of therapeutics leveraging NGF-TrkA signaling in bone can proceed.

In summary, the results from this study indicate that GA may be an attractive small molecule therapeutic to support load-induced bone formation through actions on both osteoblasts and NGF-TrkA signaling in sensory nerves. Increased skeletal adaptation is known to dramatically increase skeletal fatigue resistance^30^, so this approach may be sufficient to prevent fatigue injuries in at-risk individuals^31–34^. Although this study corroborates previous work indicating the administration of GA does not cause the same level of hyperalgesia and sensitivity induced by NGF, more study is required to determine its specific effect on skeletal sensory nerves, its dependence on endogenous osteoblastic NGF, and the mechanism of its action on osteoblasts.

## ACKNOWLEDGEMENTS

Our research is supported by the National Institute of Arthritis and Musculoskeletal and Skin Diseases and the National Institute of Dental and Craniofacial Research of the National Institutes of Health under award numbers AR074953 (RET) and DE028397 (RET). The content is solely the responsibility of the authors and does not necessarily represent the official views of the funding bodies.

---

**Supplementary Figure 1:**
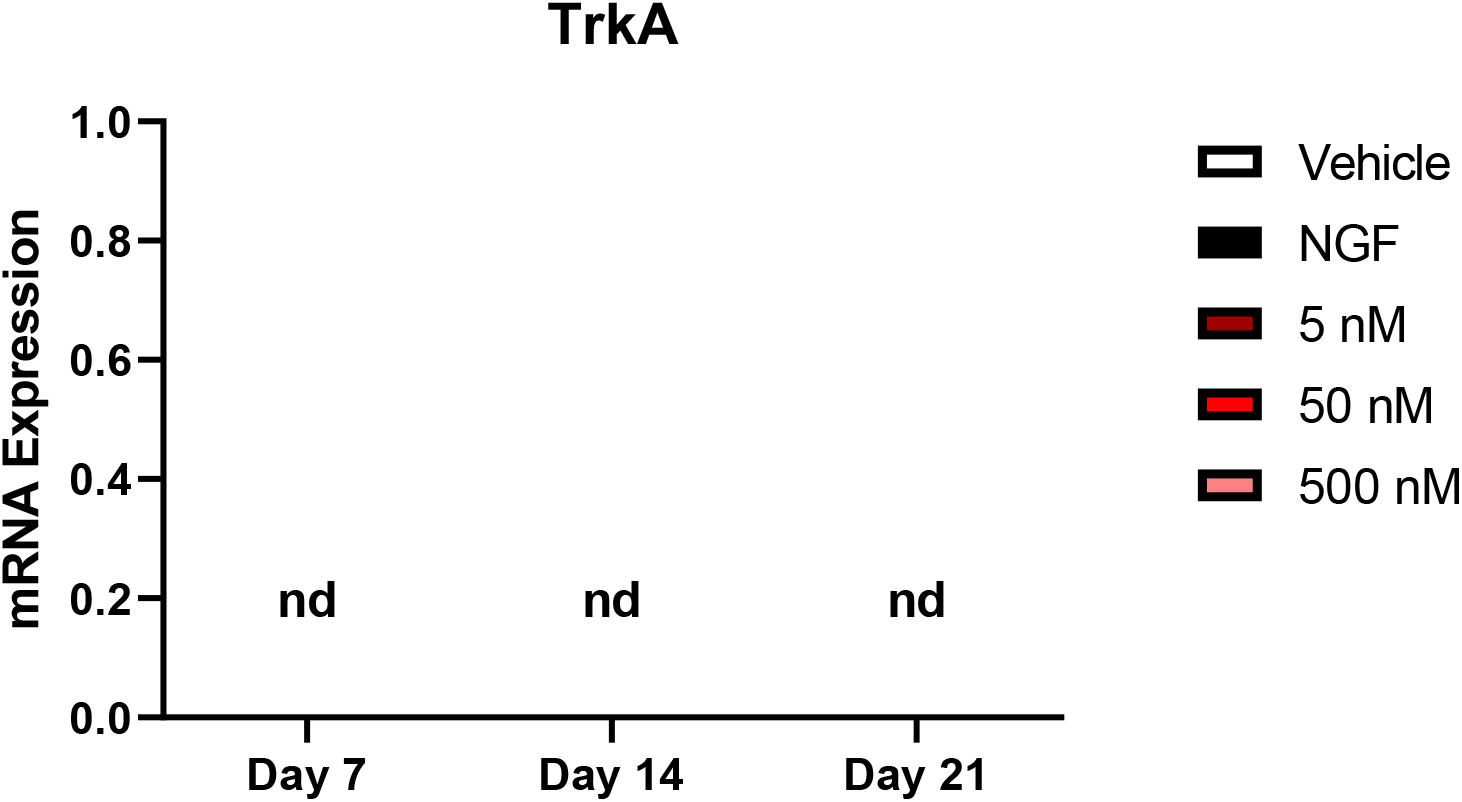
mRNA expression of TrkA in MC3T3 cells after treatment of Vehicle (DMSO), NGF, or 5, 50, 500 nM GA spanning up to 2 weeks. Nd = not detected.

## REFERENCES

1. Robling, A. G., Castillo, A. B. & Turner, C. H. Biomechanical and Molecular Regulation of Bone Remodeling. Annu. Rev. Biomed. Eng. 8, 455–498 (2006).

2. Seeman, E. Bone modeling and remodeling. Critical Reviews in Eukaryotic Gene Expression 19, 219–233 (2009).

3. Kelly, N. H., Schimenti, J. C., Ross, F. P. & van der Meulen, M. C. H. Transcriptional profiling of cortical versus cancellous bone from mechanically-loaded murine tibiae reveals differential gene expression. Bone 86, 22–29 (2016).

4. Tomlinson, R. E. et al. NGF-TrkA signaling in sensory nerves is required for skeletal adaptation to mechanical loads in mice. doi:10.1073/pnas.1701054114

5. Chermside-Scabbo, C. J. et al. Old Mice Have Less Transcriptional Activation But Similar Periosteal Cell Proliferation Compared to Young-Adult Mice in Response to in vivo Mechanical Loading. J. bone Miner. Res. Off. J. Am. Soc. Bone Miner. Res. (2020). doi:10.1002/jbmr.4031

6. Mantyh, P. W. The neurobiology of skeletal pain. Eur. J. Neurosci. 39, 508–519 (2014).

7. Castañeda-Corral, G. et al. The majority of myelinated and unmyelinated sensory nerve fibers that innervate bone express the tropomyosin receptor kinase A. Neuroscience 178, 196–207 (2011).

8. Lewin, G. R., Ritter, A. M. & Mendell, L. M. Nerve growth factor-induced hyperalgesia in the neonatal and adult rat. J. Neurosci. 13, 2136–2148 (1993).

9. Rukwied, R. et al. NGF induces non-inflammatory localized and lasting mechanical and thermal hypersensitivity in human skin. Pain 148, 407–413 (2010).

10. Apfel, S. C. Nerve growth factor for the treatment of diabetic neuropathy: What went wrong, what went right, and what does the future hold? International Review of Neurobiology 50, 393–413 (2002).

11. Bergmann, I., Reiter, R., Toyka, K. V & Koltzenburg, M. Nerve growth factor evokes hyperalgesia in mice lacking the low-affinity neurotrophin receptor p75. Neurosci. Lett. 255, 87–90 (1998).

12. Apfel, S. C. et al. Efficacy and safety of recombinant human nerve growth factor in patients with diabetic polyneuropathy: A randomized controlled trial. rhNGF Clinical Investigator Group. JAMA 284, 2215–2221 (2000).

13. Apfel, S. C. et al. Recombinant human nerve growth factor in the treatment of diabetic polyneuropathy. NGF Study Group. Neurology 51, 695–702 (1998).

14. McArthur, J. C. et al. A phase II trial of nerve growth factor for sensory neuropathy associated with HIV infection. AIDS Clinical Trials Group Team 291. Neurology 54, 1080–1088 (2000).

15. Petty, B. G. et al. The effect of systemically administered recombinant human nerve growth factor in healthy human subjects. Ann. Neurol. 36, 244–246 (1994).

16. Lee, A. C. L., Harris, J. L., Khanna, K. K. & Hong, J. H. A comprehensive review on current advances in peptide drug development and design. Int. J. Mol. Sci. 20, 1–21 (2019).

17. Lee, F. S. & Chao, M. V. Activation of Trk neurotrophin receptors in the absence of neurotrophins. Proc. Natl. Acad. Sci. U. S. A. 98, 3555–3560 (2001).

18. Jang, S.-W. et al. Gambogic amide, a selective agonist for TrkA receptor that possesses robust neurotrophic activity, prevents neuronal cell death. Proc. Natl. Acad. Sci. 104, 16329–16334 (2007).

19. Obianyo, O. & Ye, K. Novel small molecule activators of the T rk family of receptor tyrosine kinases. Biochim. Biophys. Acta - Proteins Proteomics 1834, 2213–2218 (2013).

20. Shen, J. & Yu, Q. Gambogic amide selectively upregulates TrkA expression and triggers its activation. Pharmacol. Reports 67, 217–223 (2015).

21. Longo, F. M. & Massa, S. M. Small-molecule modulation of neurotrophin receptors: a strategy for the treatment of neurological disease. Nat. Rev. Drug Discov. 12, 507–525 (2013).

22. Park, J., Fertala, A. & Tomlinson, R. E. Naproxen impairs load-induced bone formation, reduces bone toughness, and diminishes woven bone formation following stress fracture in mice. Bone 124, 22–32 (2019).

23. Hsieh, Y. L., Kan, H. W., Chiang, H., Lee, Y. C. & Hsieh, S. T. Distinct TrkA and Ret modulated negative and positive neuropathic behaviors in a mouse model of resiniferatoxin-induced small fiber neuropathy. Exp. Neurol. 300, 87–99 (2018).

24. Hens, J. R. et al. TOPGAL mice show that the canonical Wnt signaling pathway is active during bone development and growth and is activated by mechanical loading in vitro. J. bone Miner. Res. Off. J. Am. Soc. Bone Miner. Res. 20, 1103–1113 (2005).

25. Robinson, J. A. et al. Wnt/beta-catenin signaling is a normal physiological response to mechanical loading in bone. J. Biol. Chem. 281, 31720–31728 (2006).

26. Rosenbaum, T., Vidaltamayo, R., Sánchez-Soto, M. C., Zentella, A. & Hiriart, M. Pancreatic beta cells synthesize and secrete nerve growth factor. Proc. Natl. Acad. Sci. U. S. A. 95, 7784–7788 (1998).

27. Tomlinson, R. E. et al. NGF-TrkA Signaling by Sensory Nerves Coordinates the Vascularization and Ossification of Developing Endochondral Bone. Cell Rep. 16, 2723–2735 (2016).

28. Yada, M., Yamaguchi, K. & Tsuji, T. NGF stimulates differentiation of osteoblastic MC3T3-E1 cells. Biochemical and Biophysical Research Communications 205, 1187–1193 (1994).

29. Johnstone, M. R. et al. The selective TrkA agonist, gambogic amide, promotes osteoblastic differentiation and improves fracture healing in mice. 19, 1–10 (2019).

30. Warden, S. J. et al. Bone adaptation to a mechanical loading program significantly increases skeletal fatigue resistance. J. bone Miner. Res. Off. J. Am. Soc. Bone Miner. Res. 20, 809–816 (2005).

31. Milgrom, C. et al. Stress fractures in military recruits. A prospective study showing an unusually high incidence. J. Bone Joint Surg. Br. 67, 732–735 (1985).

32. Milgrom, C. et al. Youth is a risk factor for stress fracture. A study of 783 infantry recruits. J. Bone Joint Surg. Br. 76, 20–22 (1994).

33. Warren, M. P., Brooks-Gunn, J., Hamilton, L. H., Warren, L. F. & Hamilton, W. G. Scoliosis and fractures in young ballet dancers. Relation to delayed menarche and secondary amenorrhea. N. Engl. J. Med. 314, 1348–1353 (1986).

34. Johnson, A. W., Weiss, C. B. J. & Wheeler, D. L. Stress fractures of the femoral shaft in athletes--more common than expected. A new clinical test. Am. J. Sports Med. 22, 248–256 (1994).

